# GCDPipe: risk gene, cell type, and drug ranking for complex traits

**DOI:** 10.1101/2022.07.27.501775

**Authors:** Daria Pinakhina, Alexander Loboda, Alexey Sergushichev, Mykyta Artomov

## Abstract

We introduce a user-friendly machine learning tool for risk gene, cell type, and drug ranking for complex traits - GCDPipe. It uses gene-level GWAS-derived data and publicly available expression data to train a model for prediction of disease risk genes and relevant cell types. Gene-ranking information is then coupled with known drug targets data to prioritize drugs based on their estimated functional effects associated with identified risk genes. The pipeline was tested in two case studies: inflammatory bowel disease (IBD) and schizophrenia, then it was applied to Alzheimer’s disease to investigate potential options for drug repurposing. The results show that GCDPipe is an effective tool to unify genetic risk factors with cellular context and known drug targets.

## Introduction

There is a common recognition that GWAS could be a powerful tool for drug development, as well as for drug repurposing ^1,2^. The latter enables an effective transition of existing therapies into new applications in clinical practice. Several approaches to leverage GWAS data for drug discovery and repurposing have already been proposed ^1–13^.

One of the most intuitive solutions for the utilization of GWAS data in drug repurposing is to focus on the leading candidate genes identified in a GWAS as targets for drug repositioning^1^. Among the main limitations of such a strategy is the small yield of drug candidates from top associations in GWAS, as only a modest fraction (22%) of protein-coding genes is currently druggable^1^. Additional tools, such as Gentrepid^14^, use diverse information on pathway, protein–protein interaction (PPI) and/or protein domain homology to extend candidate gene lists and overcome this limitation. The other solution to candidate gene list extension is provided by machine learning-based pipelines for gene prioritization, however the utility of such methods for drug and drug-target prioritization remains understudied ^15,16^.

GWAS findings can inform development of therapeutic approaches by providing evidence for identification of disease-relevant cell types and tissues. This information could guide the development of drugs with increased cell type specificity to minimize potential side effects ^17,18^. Despite a rapid increase in the amount of scRNA-seq data, aligning GWAS results with cell-type specific expression pattern is the area of active development with various approaches already established ^20,21,22,19,11^. Most of them are based on the linkage of variant-level association signals from GWAS to cell-type-specific expression through eQTL properties. At the same time, the complexity of direct implication of disease-causal genes from GWAS variant data remains the main challenge in understanding disease biology through genetic studies, indicating that more flexible solutions without a tight linkage to variant-level statistics would be required.

Here, we introduce a random forest classifier-based tool, GCDPipe, for GWAS-derived gene-level results analysis, which allows a joint extension of a risk gene list, gene ranking by probability to influence disease risks, and prioritization of expression profiles, representing cell types and/or tissues, based on their importance for risk gene identification. The obtained gene ranking is then used to prioritize drugs using drug-gene interaction. The utility of the pipeline is illustrated in several case studies: inflammatory bowel disease (IBD) [MIM: 266600] and schizophrenia [MIM: 181500]. It is subsequently applied to Alzheimer’s disease [MIM: 104300] to prioritize potential drug candidates for repurposing.

## Material and Methods

### GCDPipe Input

#### Feature data

GCDPipe uses as input a feature matrix with expression values across cell types/tissues that can be constructed from a wide range of publicly available expression atlases (**Sup. Table S1, Fig. 1**).

**Fig. 1.**
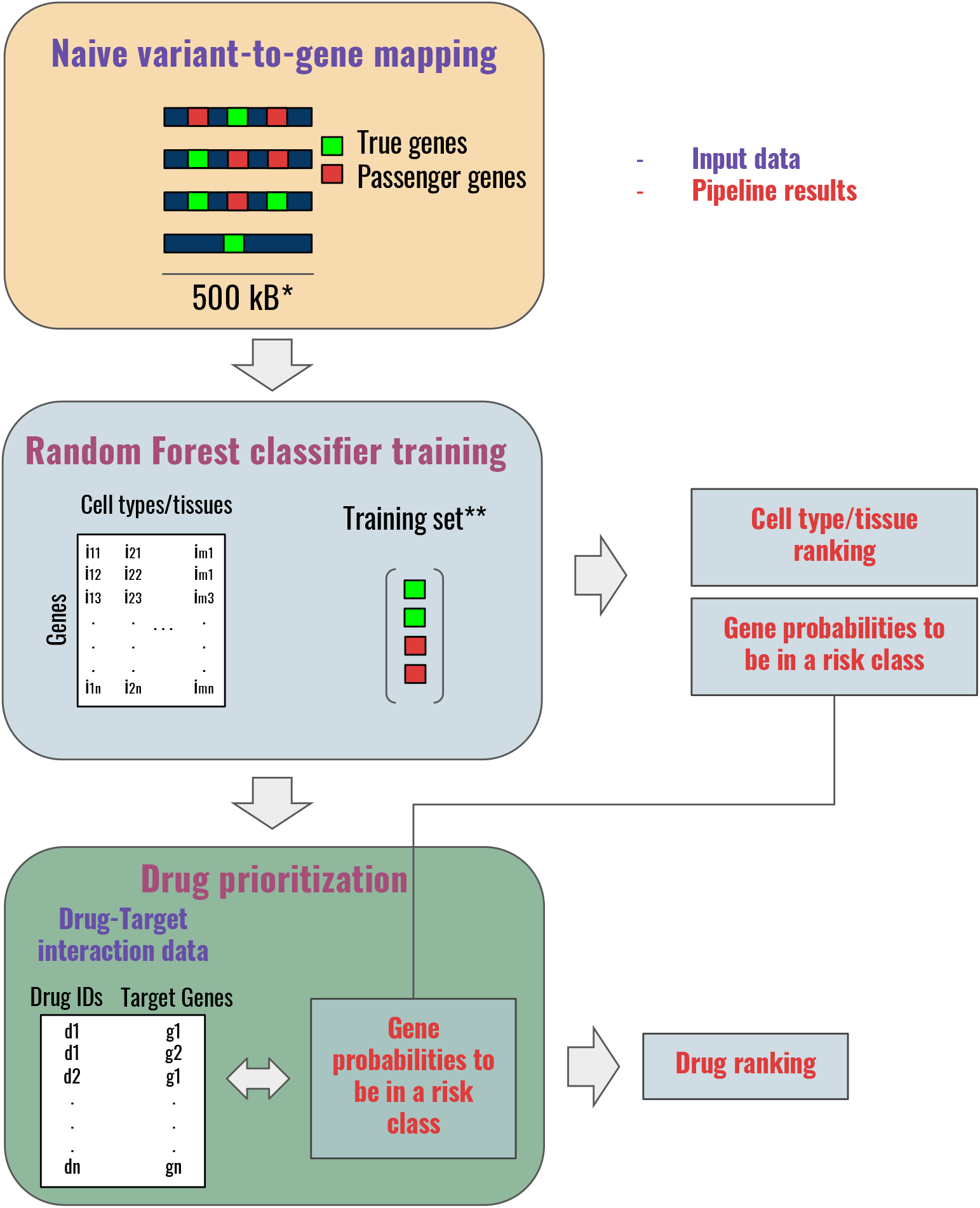
A general pipeline scheme for joint risk gene identification, cell/tissue type ranking, and drug prioritization for a disease based on GWAS variant-to-gene mapping results. *- adjustable parameter; ** - is split into training and validation sets for cross validation during hypermarameter optimization; a custom fraction of genes can also be allocated for the test set.

#### Gene data for training and testing

Next, GCDPipe requires a set of genes that are marked as ‘causal’ and ‘non-causal’ for training and testing. Alternatively, a list of GWAS loci with the ‘causal’ genes can be provided for gene set generation (**Sup. Methods**).

#### Drug-target interaction data

There is an option to run GCDPipe to return drug ranking based on maximum probability of any drug-targets to be assigned to the disease-risk class of genes. In this case, the pipeline requires a matrix with drug identifiers and their gene targets.

#### Trait-specific drug list

If a set of drugs for a phenotype of interest is known, GCDPipe can use this information to perform an initial quality assessment of gene and drug ranking. Two options are provided. First is to compare the gene scores between the gene-targets of the provided true set of drugs and all other genes. Second is to compare drug scores between the provided true set of drugs and all other drugs.

### Classifier *training and testing settings*

GCDPipe also allows to set a fraction of genes to be used for testing, and a range of hyperparameters to tune the random forest classifier (**Sup. Methods**).

We provide examples of input files for IBD and schizophrenia case studies at https://github.com/ACDBio/GCDPipe.

### GCDPipe algorithm

There are two key steps in gene classification within GCDPipe. First, a random forest classifier is trained to distinguish between risk and non-risk genes. Second, the performance of this classifier is assessed to ensure the proper quality of training. An original set of input genes is initially split into two parts - a training and a testing set. The first one is used solely for the classifier training and the other is set aside for future performance evaluation (**Sup. Fig. S1A, Fig. 1**).

Random forest classifier training involves hyperparameter tuning with GridSearchCV using five-fold cross-validation. The classifier is subsequently tested using a test gene set (**Sup. Fig. S1A**).

The ranking of expression profiles (representing cell types/tissues) is performed using a modification of the feature importance analysis with SHAP values (**Sup. Fig. S2**). The ranking score represents the correlation between SHAP values for the risk class for a feature (expression profile) and gene expression values for the same feature (**Sup. Methods**).

Next, a simplistic drug prioritization scheme is applied: given the information on drugs and their targets, the drugs are arranged by maximal probabilities of any of their gene-targets to be assigned to the risk class.

### GCDPipe output

As an output, GCDPipe returns: a range of quality metrics for the obtained classifier and its ROC curve; a matrix with gene probabilities to influence disease risk and their risk class assignment using the probability threshold corresponding to the largest difference between true positive and false positive rates; a matrix with expression profiles’ (characterizing cells/tissue types) scores.

If a drug analysis is performed, a matrix with drug scores is returned. If a drug set of interest is provided, GCDPipe can run two Mann-Whitney tests: comparing the gene scores of all targets of the given drugs with those of other genes and comparing their drug scores with scores of all other drugs.

### Case study experimental design

We used the results of variant-to-gene mapping from GWAS studies on IBD, schizophrenia, and Alzheimer’s disease^23–27^ and assembled the sets of genes for classifier training and testing based on them (**Sup. Methods**, *Input files specifics and their processing*). The feature matrices were obtained using publicly available expression datasets, and the information on drug targets was obtained from DGIDB^34^ and DrugCentral^35^ (only the drugs with DrugBank IDs were considered) (**Sup. Table S1**).

To check whether the gene ranking given by GCDPipe is consistent with other sources of evidence on disease genetic background, we performed an enrichment analysis of 500 top scoring genes with top 200 genes in differential expression datasets characterizing a range of conditions. Additionally, as a positive control for the schizophrenia case study, we used 200 genes with the lowest Q-values obtained from the schizophrenia exome sequencing project (SCHEMA)^36^ (**Sup. Table S1**).

Next, to assess the power of GCDPipe gene ranking for drug target search, we performed enrichment analyses of the gene vectors arranged by risk probabilities with targets of the drugs of 4 selected diseases (IBD, schizophrenia, coronary artery disease [MIM: 608320] and coronary heart disease [MIM: 607339] (CAD and CHD), asthma [MIM: 600807]). In addition, we compared the scores for the drug categories for the corresponding diseases with those for random drug selections and for muscle relaxants (M03 ATC code). The disease-specific drug sets were obtained from DrugBank^37^.

For the application study on Alzheimer’s disease, we compared enrichments of the gene ranking with drug targets for all ATC drug categories, including anti-dementia (N06D) drugs.

In addition, we performed KEGG^38^ pathway enrichment of 500 top ranking risk genes and of the ‘true’ genes from the classifier training and testing sets for schizophrenia to assess the applicability of the GCDPipe for uncovering disease-relevant molecular mechanisms.

We also tested the pipeline performance with varying training risk gene count for schizophrenia. It was run with 2, 5, 14, 24 risk genes; enrichment of the top ranking genes with the genes of the SCHEMA gene set and the enrichment of the gene ranking with schizophrenia drug targets were estimated as performance measures.

The risk genes used for classifier training and testing were excluded from enrichment analyses, as well as from drug target enrichments. The examples of code used in the analysis of the results are provided at https://github.com/ACDBio/GCDPipe.

## Results

### Case study 1: Inflammatory bowel disease (IBD)

27 IBD risk loci bearing likely causal genes identified in the studies by Huang et al., 2017^23^, Sazonovs et al., 2021^24^, and a range of other studies, were used to train and test GCDPipe. The list of 27 likely causal genes was assembled as follows:

11 genes were chosen due to presence of protein-coding and eQTL variants in signals mapped to less than 50 variants, at least one of which has posterior probability > 50% for IBD from the study by Huang et al., 2017 (see Table 1 in the original article^23^). *CYLD* gene, located immediately adjacent to *NOD2* in IBD susceptibility locus, was also included in the risk gene set, as it was hypothesized to regulate *NOD2* signaling^25,26^.

From the study by Sazonovs et al., 2021, we selected 9 coding variants with study-wide significance and uniquely mapped to genes (see Table 2 in the original article^24^). In addition, we also selected the gene *ATG4C*, which displayed a significant signal in gene-based rare-variant (MAF < 0.001) burden tests.

Finally, *RNF186* risk gene was selected based on the deep resequencing study by Beaudoin et al., 2013^27^, *ADCY7* - based on the study by Luo et al.^28^, *INAVA* - based on a range of studies^29,30^, *SLAMF8 and PLCG2*, which bear likely causal missense variants in the GWAS by de Lange et al., 2017^31^ were also included.

27 genes were used as a positive class and other genes within +/-250 kB window of each of the positive genes were selected as a negative class (in total 343 genes). They were randomly split as 70% training and 30% testing (**Sup. Methods**, *Input files specifics and their processing*).

The resultant risk gene classifier returned AUC-ROC of 0.82 for the test set (**Sup. Fig. S1B**). The three sets of IBD-related differential expression-based genes used in the analysis showed a significant over-representation in the top scoring genes (**Fig. 2A**).

**Fig. 2.**
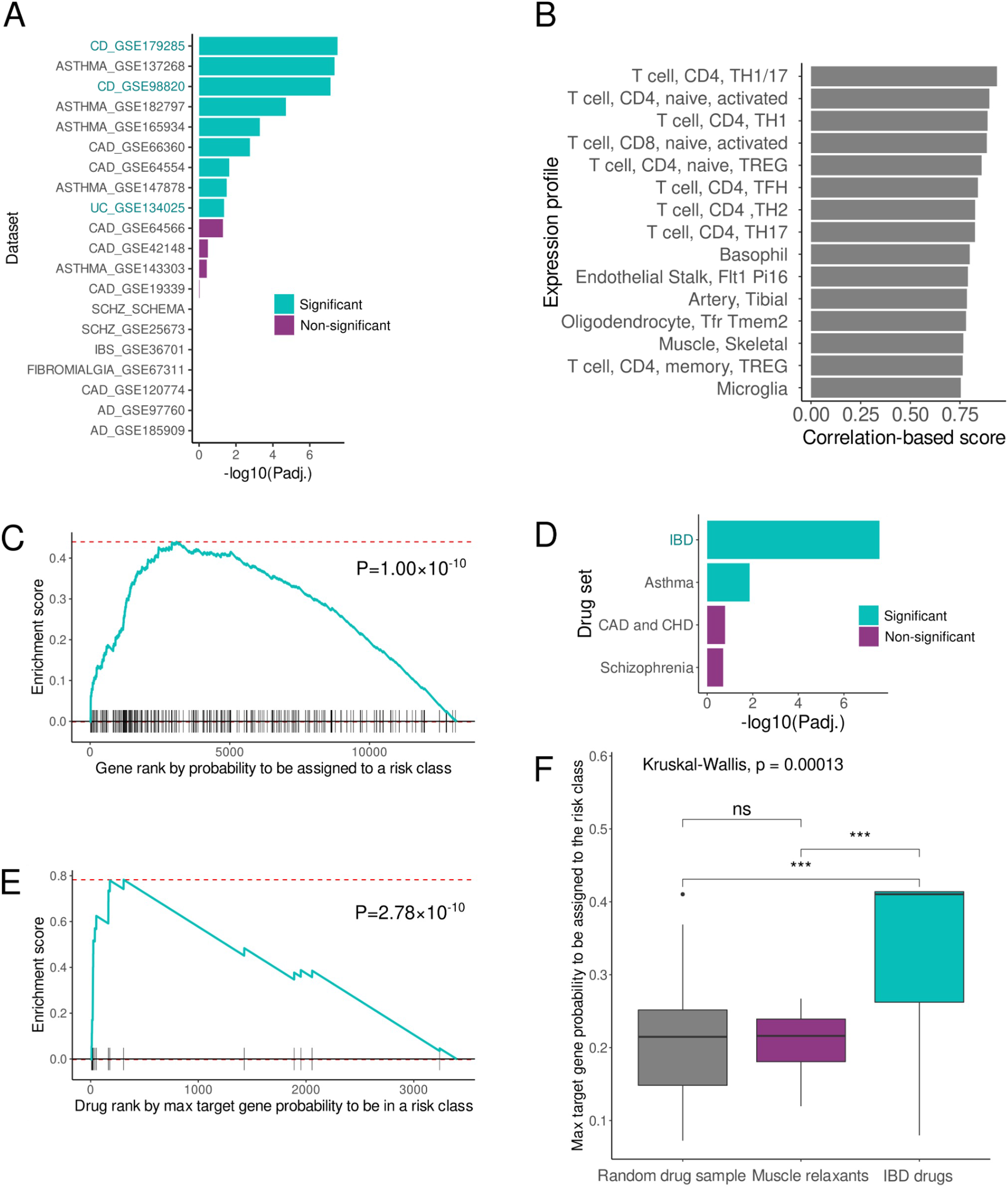
Case Study 1: IBD. **(A)** Over-representation of top 200 genes across a range of datasets among the genes* with the highest probabilities (top 500 genes) to be assigned to the risk class; **(B)** Cell types which have shown the highest correlation of their gene expression with the SHAP values for the risk class. **(C)** Enrichment of the genes* arranged by their probabilities to be assigned to the risk class with IBD drug targets. **(D)** Comparison** between significance of enrichment of the genes* arranged by their probabilities to be assigned to the risk class with drug targets across a range of diseases. **(E)** Enrichment of drugs arranged by max probabilities of their targets to be assigned to the risk class with IBD drugs. **(F)** Comparison between max target probabilities to be assigned to the risk class for IBD drugs, a random drug sample of the same size, and muscle relaxants (M03 ATC code, the control drug group). * - excluding GWAS risk genes used for classifier training. ** - The p-values are adjusted for comparison with drugs from all ATC categories and the disease-specific drug sets.

Among the 490 expression profiles used for classifier training, CD4+ T cells (TH cells, in particular), which are known major initiators of IBD, prevailed in the top 10 prioritized profiles (for CD+ T cells p=7.17×10^−11^, Fisher exact test) (**Fig. 2B**).

A significant (P_adj._=1.00×10^−10^, empirical) enrichment of the genes arranged by probabilities to be assigned to the risk class with targets of 24 IBD drugs was observed (**Fig. 2C**). Enrichment for IBD drugs was the most significant among 4 diseases used in comparison (**Fig. 2D**). IBD drugs were also significantly (P_adj._=2.78×10^−10^, empirical) enriched in the obtained drug ranking (**Fig. 2E**). The scores obtained for IBD drugs were significantly higher compared to a random set of drugs of the same size and a negative control set of muscle relaxants (p=0.00013, Mann-Whitney, two-sided) (**Fig. 2F**).

### Case study 2: Schizophrenia

GCDPipe was trained on a set of 48 loci, for which ‘true’ genes either having all variants within a 95% credible set of a genome-wide significant locus within their transcript boundaries or being single genes with Q-value in SCHEMA <=0.05 within such loci were identified by Trubetskoy et al., 2022^32^ (Supplementary Materials, Sup. Fig. S1C). In total, 121 genes were used for training and 52 - for testing. The resultant classifier showed a ROC-AUC of 0.91 on the test set. The most significant enrichment of the top scoring genes in the classifier ranking was with the leading genes from the SCHEMA exome sequencing study (P_adj._=9.23×10^−9^, hypergeometric). The top genes from the other schizophrenia-related dataset, GSE25673, also showed a significant over-representation (P_adj._=1.36×10^−3^, hypergeometric; Fig. 3A).

**Fig. 3.**
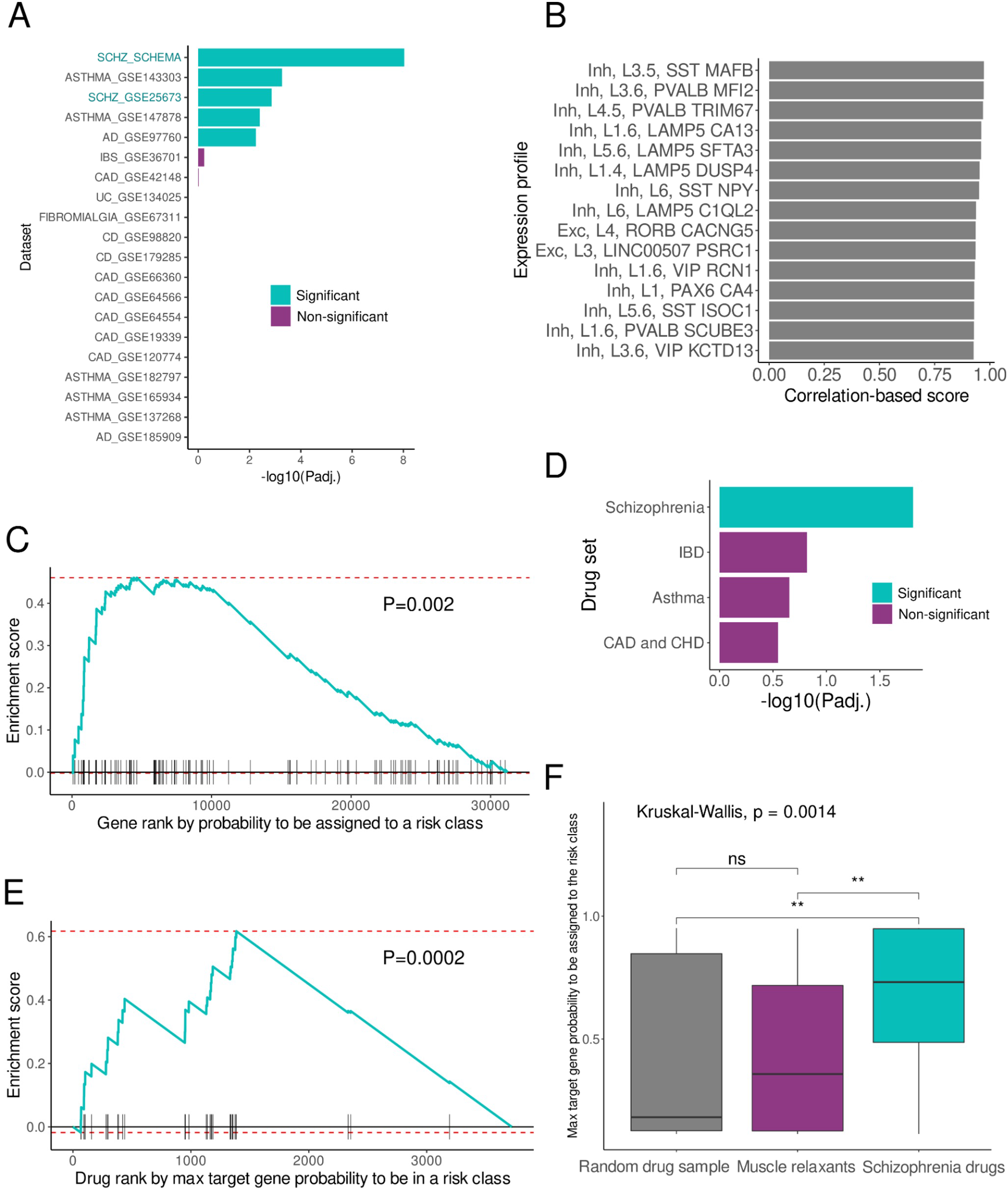
Case Study 2: Schizophrenia. **(A)** Over-representation of top 200 genes* across a range of datasets among the genes with the highest probabilities (top 500 genes) to be assigned to the risk class. **(B)** Cell types which have shown the highest correlation of their gene expression with the SHAP values for the risk class. **(C)** Enrichment of the genes* arranged by their probabilities to be assigned to the risk class with schizophrenia drug targets. **(D)** Comparison** between significance of enrichment of the genes* arranged by their probabilities to be assigned to the risk class with drug targets across a range of diseases. **(E)** Enrichment of drugs arranged by max probabilities of their targets to be assigned to the risk class with schizophrenia drugs. **(F)** Comparison between max target probabilities to be assigned to the risk class for schizophrenia drugs, a random drug sample of the same size, and muscle relaxants (M03 ATC code, the control drug group). *- excluding GWAS risk genes used for classifier training. ** - The p-values are adjusted for comparison with drugs from all ATC categories and the disease-specific drug sets.

Inhibitory neurons prevailed among the prioritized cell expression profiles (**Fig. 3B**), with subtypes^39^ of somatostatin-expressing (SST) and parvalbumin-expressing (PVALB) interneurons on the leading positions of the ranking.

The gene ranking obtained was significantly (P=0.002, empirical) enriched with schizophrenia drug targets (**Fig. 3C**), and it was the only disease for which such enrichment had been observed among the 4 conditions compared (**Fig. 3D**). We also noted a significant (P=0.0002, empirical) enrichment of the resulting drug ranking with schizophrenia drugs (**Fig. 3E**), and the scores for this drug group were significantly higher than for a random drug selection or for a control set of muscle relaxant drugs (P=0.0014, Mann-Whitney, two-sided) (**Fig. 3F**).

The top significantly enriched KEGG pathways among the 500 best scoring risk genes, excluding those used for training and testing, were dopaminergic synapse (P_adj._=1.00×10^−11^, hypergeomtetric), retrograde endocannabinoid signaling (P_adj._=6.59×10^−10^, hypergeomtetric), nicotine addiction (P_adj._=2.38×10^−9^, hypergeomtetric) and MAPK signaling pathway (P_adj._=4.22×10^−9^, hypergeomtetric) (overall, there were 71 significantly enriched pathways), while the only enriched pathway for risk genes used for training and testing was GnRH secretion (P_adj._=4.54×10^−5^, hypergeomtetric) (**Fig. S3**).

We tested parameter sensitivity to show that enrichment with SCHEMA genes and with schizophrenia drug targets is robust with respect to the size of training gene set (**Fig. S4**).

### Application study: Alzheimer’s disease

Training and testing GCDPipe using 16 Alzheimer’s disease risk loci with ‘true’ genes identified as having the scores in gene evidence rankings more than 1.5 times higher than those of other genes within a locus based on the study by Schwartzentruber et al., 2021^33^, had AUC-ROC of 0.82 on the test set (**Fig. 4A**). In total, 164 genes were used for training and 71 - for testing. A significant enrichment of the top resulting risk genes was observed with the most differentially expressed genes from the GSE97760 Alzhemer’s disease study, whereas no overlap was detected with another disease-specific dataset (GSE97760) (**Fig. 4B**).

**Fig. 4.**
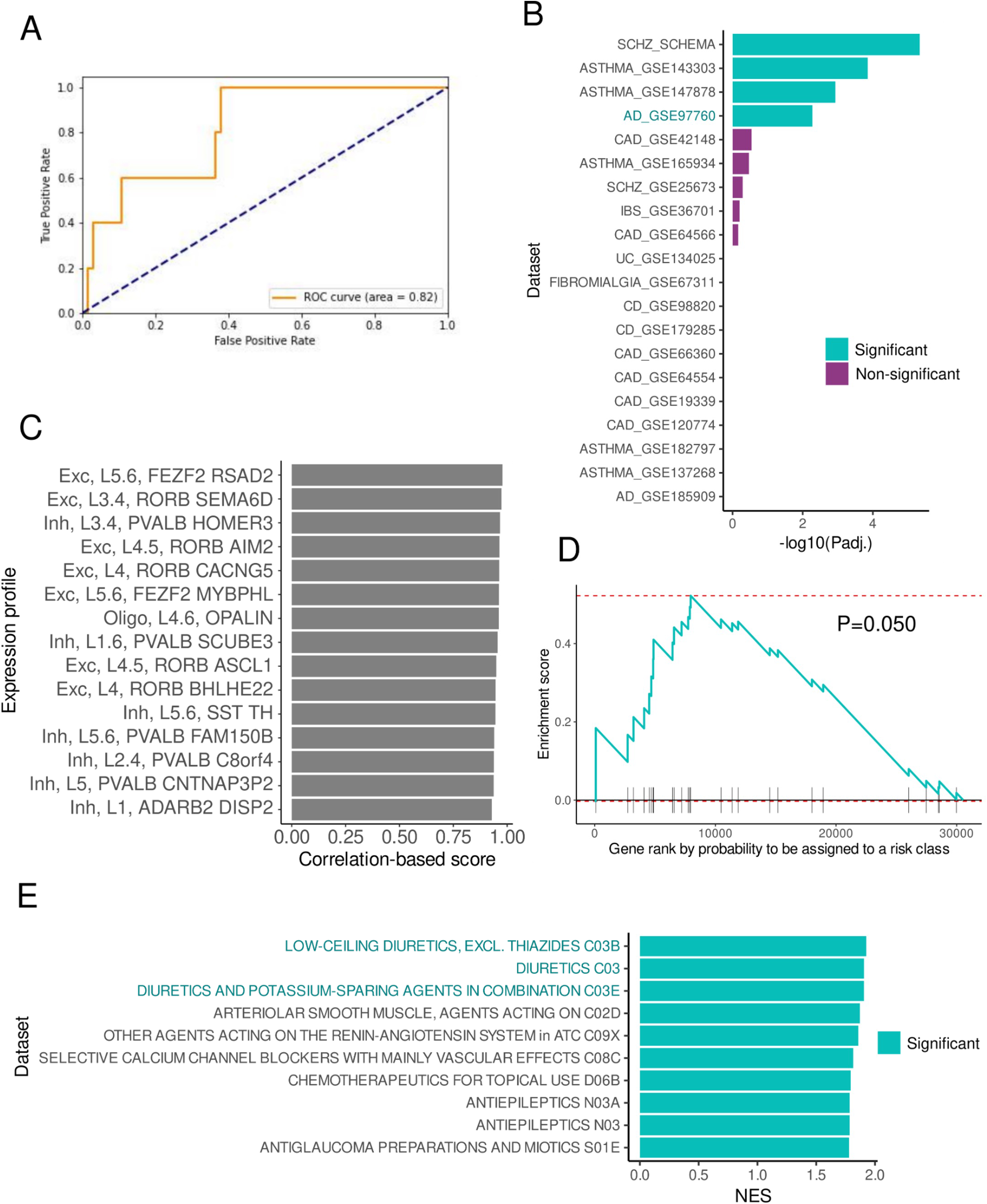
Application case: Alzheimer’s disease. **(A)** ROC curves and AUC-ROC for the obtained classifiers. (B) Over-representation of top 200 genes* across a range of datasets among the genes with the highest probabilities (top 500 genes) to be assigned to the risk class. **(C)** Cell types which have shown the highest correlation of their gene expression with the SHAP values for the risk class. **(D)** Enrichment of the genes* arranged by their probabilities to be assigned to the risk class with anti-dementia (N06D) drug targets. **(E)** Top 10 ATC drug categories, displaying maximal NES scores for enrichment of their targets in the vector of genes*, arranged by probabilities to be assigned to the risk class. *- excluding GWAS risk genes used for classifier training.

The prioritized expression profiles comprised both excitatory and inhibitory neurons, and an oligodendrocyte subtype (**Fig. 4C**).

We observed a tendency (P=0.050, Mann-Whitney, two-sided) towards enrichment with anti-dementia (N06D) drug targets of the obtained gene ranking (**Fig. 4D**). The drug categories, which display the highest NES scores for their target enrichment, were three groups of diuretics (**Fig. 4E**).

## Discussion

The results of the pipeline testing indicate that current advancements in variant-to-gene mapping for GWAS provide sufficient gene-level data to train an algorithm that achieves good risk gene discrimination on test sets even in supervised training context in case of complex traits. For both case studies, gene classification results align with independently acquired data on disease genetic architecture using differential expression and exome sequencing analyses. At the same time, the resultant gene ranking is enriched with known drug targets; and drug ranking is enriched with drugs for the corresponding conditions.

For both case studies, a disease-relevant cell type prioritization was observed. For IBD, CD4+ T cells are the leading cell types with TH17, TH1/17, and TH1 among the top prioritized subtypes. CD4+ T helper cells are considered to be the main initiators of IBD: They are enriched in the lesional tissues of patients with IBD, while their blockade or depletion is an effective treatment strategy^40^. TH17 CD4+ cells are reported to be especially important for IBD development, and in the mouse model of TH17-driven IBD, transition of TH17 precursors to TH1-like cells is shown to be absolutely required for the disease. It has been suggested that focusing on TH17 and TH1-like cells may be a promising avenue for therapy design^41^.

Markers of PVALB and SST interneurons, which were found among the leading cell types for schizophrenia, are strongly associated with regional changes in functional MRI variability, while the genetic risk of schizophrenia is shown to be enriched among genes linked to interneurons, predicting the amplitude of the cortical signal in regions dominated by PVALB. Selective modulation of GABAergic system is proposed as a promising intervention for the treatment of schizophrenia-associated cognitive defects^42,43^.

Interestingly, the leading enriched drug target categories for Alzheimer’s disease appeared to be diuretics. The use of diuretics has been shown to be associated with a reduced risk of Alzheimer’s disease, and diverse experimental and real-world evidence appeared supporting the computational repurposing of diuretic bumetanide for the treatment of the disease^44-46^. GCDPipe links GWAS-derived genetic data on the disease with these observations.

The leading enriched pathway in top ranking genes for schizophrenia is dopaminergic synapse, whereas alterations in dopaminergic neurotransmission are considered as one of the most robust pathophysiological observations in schizophrenia patients^47,48^. Another contributor to schizophrenia pathogenesis, proposed as a prospective target for its treatment and measurement, is the endocannabinoid system, and retrograde endocannabinoid signaling is the second most enriched pathway in the top ranking genes^49–53^. Nicotine addiction is the third most enriched KEGG gene set, and it complies with the observation of an association between schizophrenia and tobacco smoking^54^. These pathways are interconnected with other enriched gene sets, including glutamatergic synapse and calcium signaling. Further analysis of the resultant gene-pathway network could reveal how these mechanisms are interconnected at the molecular level. None of these pathways is enriched in the GWAS-derived risk gene set used for model development: GCDPipe converges genetic risk factors for complex traits with findings on their pathophysiology at other hierarchical levels.

## Supplemental Information

Supplemental information includes supplemental methods (a note on GCDPipe implementation and testing procedure), three figures and one table.

## Data and code availability

The code, installation and usage instructions, examples, and input data described in the case studies are available at https://github.com/ACDBio/GCDPipe.

