## Supplemental Information for "GCDPipe: risk gene, cell type, and drug ranking for complex traits"

**Supplemental Methods: GCDPipe implementation and testing** **2**

**Figure S1. Classifier testing procedure.** **5**

**Figure S2. Expression profile importance-based scoring scheme implemented in the pipeline.** **6**

**Figure S3. Results of KEGG pathway enrichment analysis for the schizophrenia case study.** **8**

**Figure S4. Evaluation of pipeline performance with varying training risk genes based on schizophrenia data.** **8**

**Table. S1. The data used for each step during pipeline testing and application.** **9**

**Supplemental References** **10**

### Supplemental Methods: GCDPipe implementation and testing

*Input files specifics and their processing*

If the input gene data is represented by loci with coordinates of the leading variants in GRCh38 and risk ('True') genes specified (in such a file, the fields "rsid", "location", "chromosome", "pipe_genesymbol" are expected; if rs id is unknown, the field can be left blank), gene coordinates from gencode v39[1] are used to subset all genes within +/- 250 kb windows of the leading variants marking them as "False", except for the specified risk genes. In such a manner, a dataset of train and test genes is obtained when it is not provided directly . Center coordinates of the loci may be not tied to variants, in this case the "rsid" field can be just left blank. Another type of input gene data supported - a file with direct specification of "True" and "False" genes (the fields 'pipe_genesymbol' and 'is_True' are expected). A dataset of features - expression profiles is supplied to the pipeline as a separate .csv file. Presence of a "pipe_genesymbol" column is expected with gene symbols and the columns can have informative names, for example, specifying cell types/tissues. These names will be present in the output feature ranking. Normalization and scaling is not obligatory for gene classification, as the pipeline is based on the Random Forest classification algorithm.

Another possible input file can be provided, containing 2 columns - one specifying Drugbank ID or drug ID in other nomenclature (the field should be named "DRUGBANK_ID") and the other - gene symbols ("pipe_genesymbol"). This information is needed for drug ranking. In our case studies we used a database assembled from DGIdb[2] and DrugCentral[3] with the drugs, for which Drugbank ID was present (this data can be found at https://github.com/ACDBio/GCDPipe).

Finally, a list of drug IDs for a disease of interest can be provided to make an initial assessment of the quality of the obtained gene and drug ranking (a single "DRUGBANK_ID" field is required in this case): the pipeline will perform comparison between probabilities of the drugs' targets to be assigned to a class of risk with those of other genes as well as of the scores for selected drugs compared with all other drugs. Drug score in the pipeline is computed as maximal probability among the drugs' targets to be assigned to a risk class. Mann-Whitney U test is performed, and the pipeline gives p-values and U statistics for the tests. The test was implemented using the Scipy[4] library.

If the option to unify gene nomenclature is selected, the gene symbols from the fields "pipe_genesybol" are mapped to Human gene ontology using the data from HGNC dataset [5]. Numeric data is summarized gene-wise with the "max" function.

Within the pipeline, a dataset is generated with features and information on attribution of a gene to a risk class. A fraction of genes to be used in a test set can be specified as an input to the pipeline (in the case and application studies described in the article, we used 30% of genes for the test sets). The dataset is splitted on training and testing. Within the pipeline, pandas[6,7] and numpy[8] libraries are used for data manipulation.

*Pipeline algorithm specifics*

We used the Scikit-learn[9] library for implementation of the machine learning part of the pipeline. A Random Forest classifier is trained on the training set, with hyperparameter tuning using GridSearchCV (we used the pactools[10] modification of GridSearchCV with progress bar for displaying progress in hyperparameter search). Search space for a number of hyperparameters can be changed within the pipeline. In our case studies number of estimators was set to 3000, max tree depth - at 1, min number of samples per leaf - [5,10,15,20,25,30], min number of samples for split - [2,5,10,20,30,40], max features - ['auto', 'sqrt']. In the Dash interface the latter parameters are set to a single value (to make classifier training faster), but up to 10 values can be set for them.

SHAP[11] library was used then for feature (cell type/tissue) expression profile ranking. Shapley values were obtained for the class of risk and their correlation with gene expression values. Such procedure allows to prioritize such expression profiles, higher expression levels in which result in higher gene probabilities to be assigned to the class of risk. Such modification of feature importance analysis prevents prioritization of features, lower values in which favor risk class assignment as well as of the features with more complex, non-monotonous relationship between expression and risk probability.

*Note on the pipeline output*

GCDPipe reports a range of classifier performance metrics based on a testing procedure. These include precision, recall, F1 score for each of the two classes and globally, AUC-ROC. In the output gene ranking, assigned to a risk class using a probability threshold, corresponding to maximal difference between true positive and false positive rates, are marked as ‘true’. The drugs, which have at least one gene-target assigned to a risk class with this threshold are also marked as ‘true’ in the output drug ranking.

Pipeline interface was implemented with plotly Dash[12] in Python[13].

*Note on the pipeline result analysis*

For enrichment (over-representation) analyses of top 500 genes ranked by GCDpipe with top 200 genes from differential expression datasets, the genes for the latter were ranked by -log10(Padj.)*abs(LogFC). For pathway enrichment, top 500 genes with highest probabilities to be attributed to the risk class by the pipeline were selected. The analyses and visualization of enrichment results were performed with R[14] in RStudio[15,16] using the libraries fgsea[17], dplyr[18], tidyverse[19], ggplot2[20], enrichplot[21], clusterProfiler[22], ggpubr[53].

We provide code examples used for analysis of the pipeline output at https://github.com/ACDBio/GCDPipe. These include whole result post-processing procedure for the IBD case study and pathway enrichment procedure for schizophrenia.

*Notes on risk gene definition in the case and application studies*

For IBD, we used the results of the studies by Huang et al., 2017[23] and Sazonovs et al., 2021[24] to construct a set of 27 loci with corresponding causal genes.

For schizophrenia, we constructed a set of genes for training and testing the classifier by selecting 48 risk genes from the results presented in the study by PGC[25]. Genes, included in the 'true' sets were the genes, into transcript boundaries of which fell all variants within a 95% credible set (and clump has <k3.5, the parameter k3.5singleGene was equal to 1) or which had Q-value in SCHEMA <=0.05.

To construct a training-testing gene set for Alzheimer's disease we used locus-wise causal gene assessment from the study by Schwartzentruber et al., 2021[26]. The genes with the total scores in gene evidence rankings, provided by the authors, more than 1.5 times higher than those for other genes within a locus were selected as 'true' risk genes (there were 16 such genes).

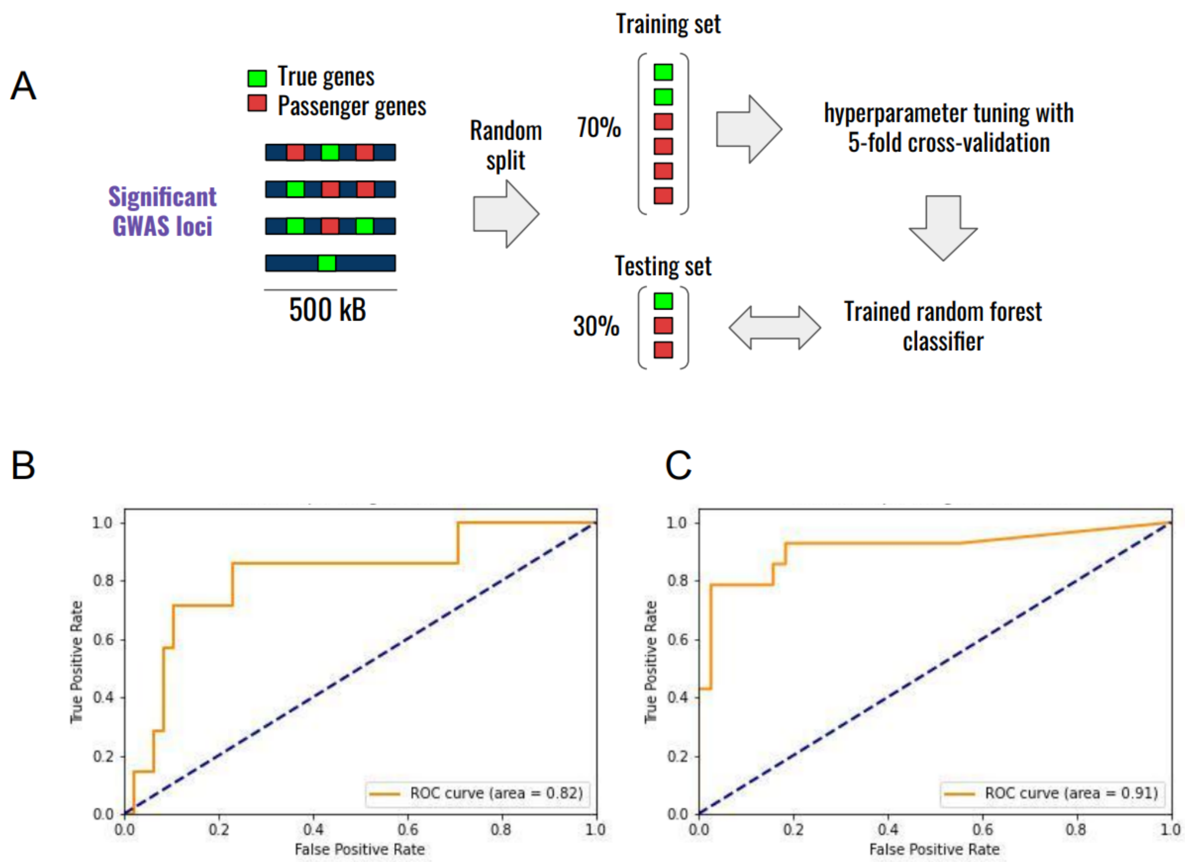


**Figure S1. Classifier testing procedure.** Training and testing datasets splitting approach (A). ROC curves and AUC-ROC for the obtained classifiers: IBD (B), Schizophrenia (C).

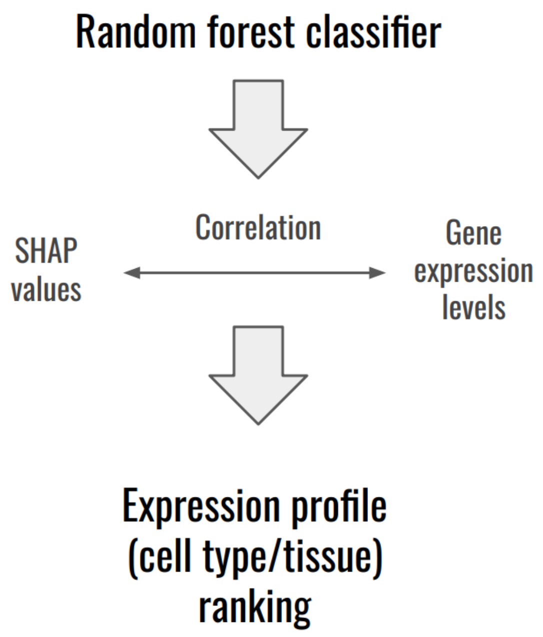


**Figure S2. Expression profile importance-based scoring scheme implemented in the pipeline.**

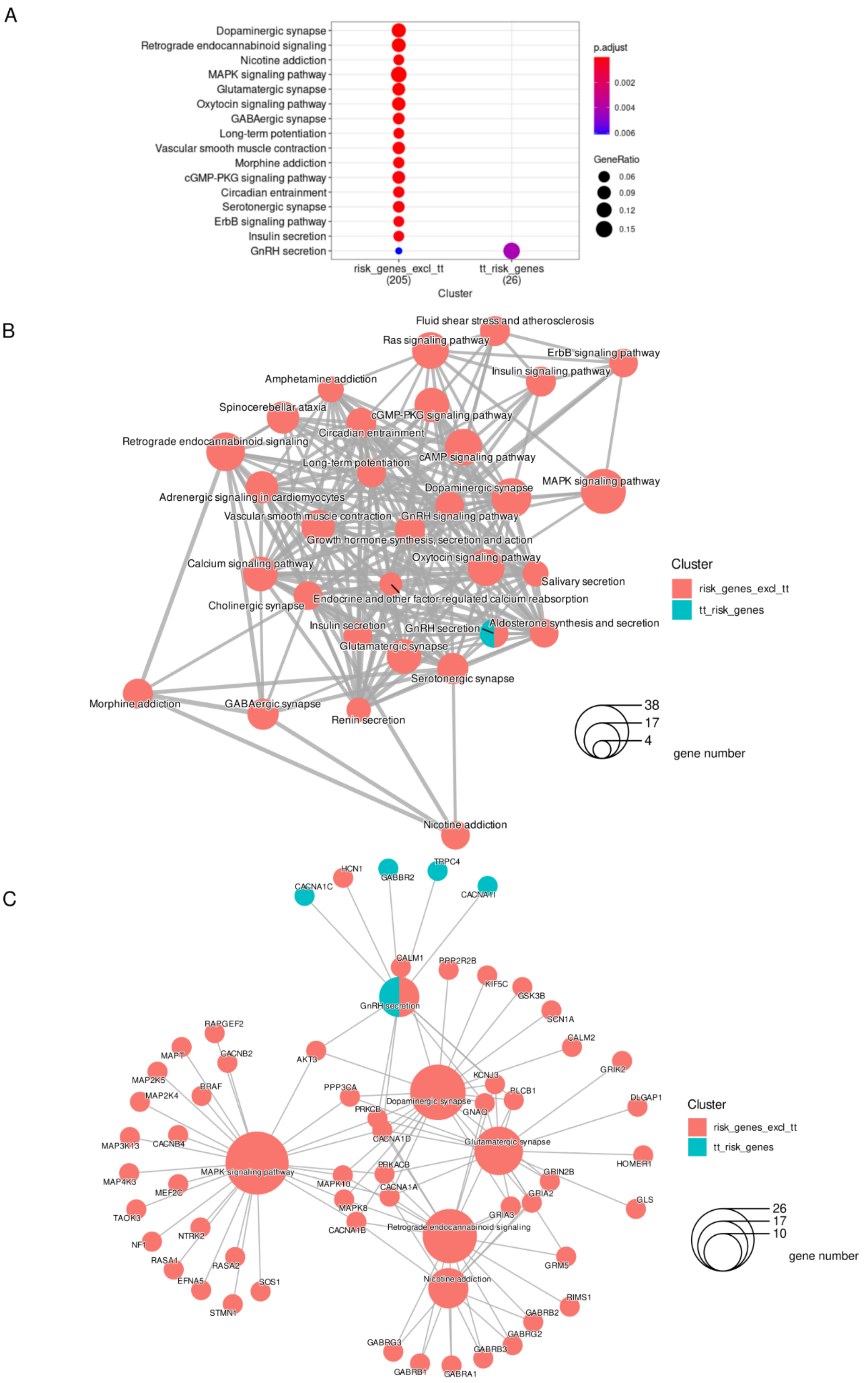
A
Dopaminergic synapse
Retrograde endocannabinoid signaling
Nicotine addiction
MAPK signaling pathway
Glutamatergic synapse
Oxytocin signaling pathway
GABAergic synapse
Long-term potentiation
Vascular smooth muscle contraction
Morphine addiction
cGMP-PKG signaling pathway
Circadian entrainment
Serotonergic synapse
ErbB signaling pathway
Insulin secretion
GnRH secretion
risk_genes_excl_tt
(205)
tt_risk_genes
(26)
Cluster
p.adjust
0.002
0.004
0.006
GeneRatio
0.06
0.09
0.12
0.15
B
Fluid shear stress and atherosclerosis
Ras signaling pathway
ErbB signaling pathway
Insulin signaling pathway
Amphetamine addiction
Spinocerebellar ataxia
cGMP-PKG signaling pathway
Circadian entrainment
Retrograde endocannabinoid signaling
cAMP signaling pathway
Long-term potentiation
MAPK signaling pathway
Dopaminergic synapse
Adrenergic signaling in cardiomyocytes
Vascular smooth muscle contraction
GnRH signaling pathway
Growth hormone synthesis, secretion and action
Oxytocin signaling pathway
Calcium signaling pathway
Salivary secretion
Endocrine and other factor-regulated calcium reabsorption
Cholinergic synapse
Aldosterone synthesis and secretion
Insulin secretion
GnRH secretion
Glutamatergic synapse
Serotonergic synapse
Morphine addiction
GABAergic synapse
Renin secretion
Nicotine addiction
Cluster
risk_genes_excl_tt
tt_risk_genes
38
17
4
gene number
C
HCN1
TRPC4
GABBR2
CACNA1C
CACNA1I
CALM1
PPP2R2B
KIF5C
GSK3B
GnRH secretion
SCN1A
RAPGEF2
CALM2
MAPT
CACNB2
AKT3
GRIK2
MAP2K5
KCNJ3
BRAF
PPP3CA
PLCB1
DLGAP1
MAP2K4
Dopaminergic synapse
PRKCB
GNAQ
MAP3K13
CACNB4
CACNA1D
Glutamatergic synapse
HOMER1
PRKACB
MAP4K3
MAPK signaling pathway
GRIN2B
MEF2C
MAPK10
CACNA1A
GRIA2
GLS
TAOK3
MAPK8
GRIA3
NTRK2
CACNA1B
Retrograde endocannabinoid signaling
NF1
RASA1
RASA2
GRM5
SOS1
EFNA5
Nicotine addiction
STMN1
RIMS1
GABRB2
GABRG2
GABRG3
GABRB3
GABRB1
GABRA1
Cluster
risk_genes_excl_tt
tt_risk_genes
26
17
10
gene number

**Figure S3. Results of KEGG pathway enrichment analysis for the schizophrenia case study.** Dotplot, comparing the leading 15 significantly enriched pathways in top 500 genes identified with GCDPipe, excluding 'true' GWAS-derived risk genes and in the set of 'true' genes used for training and testing **(A)**. Enrichment map, showing a network representation of the pathways over-represented in the two gene sets (the stronger the similarity, the shorter and thicker the edges) **(B)**. Linkage gene-pathway network for 5 pathways most significantly enriched in the two gene sets **(C)**.
risk_genes_excl_tt - gene set of top 500 genes with maximal probabilities to be assigned to the risk class, excluding GWAS-derived risk genes used for classifier training and testing;
tt_risk_genes - gene set of GWAS-derived risk genes used for classifier training and testing

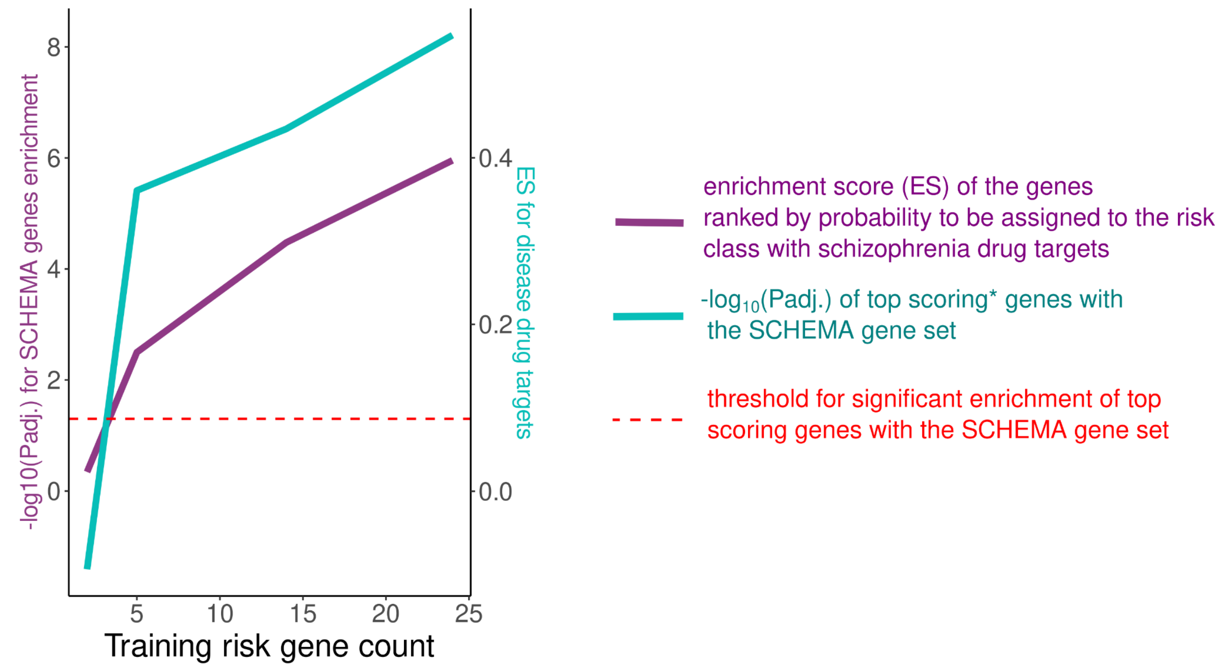


**Figure S4. Evaluation of pipeline performance with varying training risk genes based on schizophrenia data.** *top 500 genes by probability to be assigned to the risk class

| Disease | GWAS genetic fine-mapping results source | Sources of expression data used for classifier training | Datasets used in the gene prioritization results' validation | Drug-target genes interaction data sources | Disease drug sets |
|---|---|---|---|---|---|
| IBD | Huang et al.,2017[23] Sazonovs et al., 2021 (preprint)[24] | DICE Immune Cell Atlas[27], GSE125970 (Epithelial Intestinal Cells)[28], Human Lung Cell Atlas (Blood Cells)[29], GTEx[30], DropViz[31] | SCHEMA Exome sequencing data (SCHZ)[32], GSE25673 (SCHZ)[33], GSE179285 (CD)[34], GSE98820(CD)[35], GSE134025 (UC)[36], GSE36701 (IBS)[37], GSE143303 (ASTHMA)[38], GSE147878 (ASTHMA)[39], GSE182797 (ASTHMA)[40], GSE165934 (ASTHMA)[41], GSE137268 (ASTHMA)[42], GSE97760 (AD)[43], GSE185909 (AD)[44], GSE66360 (CAD)[52], GSE64566 (CAD)[45], GSE64554 (CAD)[45], GSE19339 (CAD)[46], GSE120774 (CAD)[47], GSE67311 (FMS)[48] | DGIdb[2], DrugCentral[3] | DrugBank[49] |
| Schizophrenia | PGC, 2022[50] | Human Multiple Cortical Areas SMART-seq (Allen Brain Map)[51], GTEx[30] | | | |
| Alzheimer's disease | Schwartzentruber et al., 2021[26] | Human Multiple Cortical Areas SMART-seq (Allen Brain Map)[51], DICE Immune Cell Atlas[27], GTEx[30] | | | |

**Table. S1. The data used for each step during pipeline testing and application.**
